# Systematic over-expression screens for chromosome instability identify conserved dosage chromosome instability genes in yeast and human tumors

**DOI:** 10.1101/038489

**Authors:** Supipi Duffy, Hok Khim Fam, Yikan Wang, Erin B. Styles, Jung-Huyn Kim, J. Sidney Ang, Tejomayee Singh, Vladimir Larionov, Sohrab Shah, Brenda J. Andrews, Cornelius F. Boerkoel, Phillip Hieter

**Affiliations:** Michael Smith Laboratories, University of British Columbia, Vancouver, British Columbia, Canada; Child and Family Research Institute, University of British Columbia, Vancouver, Canada; Department of Medical Genetics, University of British Columbia, Vancouver, Canada; BC Cancer Agency, Vancouver, British Columbia, Canada; Department of Molecular Genetics, University of Toronto, Canada; The Donnelly Centre for Cellular and Biomolecular Research, University of Toronto, Canada; Center for Cancer Research, National Cancer Institute, Bethesda, MD, USA

**Keywords:** Dosage chromosome instability, Synthetic dosage lethality, Tdp1, Rhabdomyosarcoma, over-expression, chromosome instability

## Abstract

Somatic copy number amplifications (SCNAs) and gene over-expression are common features of many cancers. To determine the role of gene over-expression on genome stability, we performed functional genomic screens in the budding yeast for chromosome instability, a defining characteristic of cancer that can be targeted by therapeutics. Over-expression of 245 yeast genes increases chromosome instability by influencing processes such as chromosome segregation and DNA damage repair. Testing candidate human homologs, which were highly recurrently altered in tumors lead to the identification of 2 genes, Tdp1 and Taf12 that contribute to CIN in human cells when over-expressed. Rhabdomyosarcoma lines with higher levels of Tdp1 also show chromosome instability and can be partially rescued by siRNA-mediated knockdown of Tdp1. Using synthetic dosage lethality screens in yeast, we identified candidate target genes that will specifically target tumors with high levels of Tdp1. We demonstrate the utility of functional genetic screens in model organisms to broaden the spectrum of CIN genes, to identify novel genes relevant to chromosome instability in humans and to identify candidate gene targets that can be leveraged to selectively kill tumors over-expressing specific genes.

## Introduction

Somatic copy number amplifications (SCNAs) are the most prevalent genetic alterations in cancer genomes (Beroukhim et al., 2010). The increased frequency of recurring SCNAs suggests that some SCNAs may be cancer drivers and emphasizes the need to uncover driver genes within these regions (Sanchez-Garcia et al., 2014). However, as amplified regions often encompass multiple genes, defining potential driver genes on SCNAs and distinguishing driver SCNAs events still remain a challenge (Vogelstein et al., 2013; Zack et al., 2013). Remarkably, a majority (>70%) of recurrently altered regions in tumour genomes also do not contain known oncogenes or tumor suppressors (Zack et al., 2013). While it is clear that amplifications of oncogenes can drive tumor progression, it is less clear how amplifications may affect genome stability.

Genome instability is an inherent enabling characteristic of cancer and is essential for tumor initiation and progression (Hanahan and Weinberg, 2011). Alterations that cause genome instability happen early during tumour formation and promote the accumulation of errors during DNA replication, repair and chromosome segregation thereby increasing the likelihood that a cell will acquire multiple genetic changes necessary for tumor progression (Stratton et al., 2009). Chromosome instability (CIN), the most prevalent form of genome instability results in aneuploidy and is observed in a majority of tumors (Bayani et al., 2007; Lengauer et al., 1997). CIN is possibly the major contributor to intratumoral heterogeneity, the presence of genetically distinct populations of cells within a single tumor, impacting treatment strategy, drug resistance and tumour evolution (Dewhurst et al., 2014; Fisher et al., 2013; Horswell et al., 2013; McBride et al., 2015). For these reasons, defining genes and pathways that drive genome instability and understanding the mechanisms affect genome stability is extremely relevant for understanding for tumor etiology and progression. While many mutations are known to cause genome instability (Stirling et al., 2011; Yuen et al., 2007), less is known about the spectrum of genes that when amplified or overexpressed cause genome instability.

Given chromosome instability is an enabling characteristic of cancer (Hanahan and Weinberg, 2011), the use of aneuploidy to selectively target tumors with CIN have been explored as a therapeutic approach (Janssen et al., 2011; Manchado and Malumbres, 2011; Tang et al., 2011). Similarly, the alterations underlying CIN could be targeted using a synthetic lethality-based approach. For example, the synthetic lethal (SL) interaction between mutations in the genome stability gene BRCA1 and inhibition of PARP (Ashworth, 2008; Farmer et al., 2005) has been exploited to treat ovarian tumors. The concept of synthetic lethality has been a widely adapted therapeutic strategy to selectively target the tumor cells (Fece de la Cruz et al., 2015; McLornan et al., 2014; van Pel et al., 2013). Most SL approaches have focused on exploiting specific mutations. However, there are just as many amplified regions as deleted regions in cancer genomes (Zack et al., 2013) and the cumulative gene dosage balance model suggest that both amplifications and deletions are equally important determinants that drive tumor progression (Davoli et al., 2013). Thus we propose synthetic dosage lethality (SDL), which is synthetic lethality with an amplified or overexpressed gene, as an approach to selectively target tumors that over-express particular genes. Synthetic dosage lethality occurs when the over-expression of a gene is not lethal in a wild type background, but in conjunction with a second site non-lethal mutation causes lethality (Kroll et al., 1996; Measday et al., 2005; Measday and Hieter, 2002). Given the high frequency of genes amplified or over-expressed in tumours, SDL affords an approach to greatly expand the number of tumours that can be treated with a synthetic lethal-based approach.

Here we report a cross-species approach to identify genes that when over-expressed result in CIN and identify synthetic lethal partners for one such over-expressed dosage CIN genes, Tdp1. Genome-wide screens in yeast for over-expression uncovered 245 dosage CIN (dCIN) genes, 237 of which are novel. We assayed dosage CIN in a subset of human homologs of yeast dCIN genes that are recurrently amplified and/or over-expressed in tumors and found that 4 of 20 candidate human homologs recapitulated a dCIN phenotype in human cells. One of the conserved human dCIN genes Tdp1 is over-expressed in rhabdomyosarcoma cells that exhibit chromosome instability, which was reduced upon siRNA-mediated knockdown of Tdp1. We then screened yeast *TDP1* over-expression for synthetic dosage lethality and found several SDL genetic interactions, which identify potential therapeutic targets for tumors over-expressing Tdp1. Taken together we demonstrate that dCIN genes could be utilized to selectively target cancers.

## Results

### Systematic gene over-expression in yeast identifies 245 dosage CIN genes

To discover genes whose increased expression leads to genome instability, we performed two whole-genome chromosome instability screens in yeast. We used an arrayed collection of ~5200 yeast strains, each conditionally over-expressing a unique gene under the galactose promoter (Douglas et al., 2012). The Chromosome Transmission Fidelity (CTF) assay, monitors the inheritance of an artificial chromosome fragment (CF) (Douglas et al., 2012; Yuen et al., 2007) and the A-Like Faker screen (ALF) measures the loss of the *MATα* locus, thereby allowing haploid cells to mate with a tester strain of the same mating type (Strathern et al., 1981) (Figure 1A). For the CTF screen, the marker chromosome was introduced into the array using synthetic genetic array (SGA) technology (Tong et al., 2001). Overexpression was induced by plating on media containing galactose, after which the cells were placed on media with low adenine for the colony sectoring assay (Figure S1A) (Hieter et al., 1985; Spencer et al., 1990). The inheritance of the *MATα* locus was monitored for the ALF assay, using a mating test following gene induction on galactose (Figure S1B). Genes identified in the genome-wide screens were subjected to additional validations. First, genes identified from the primary screens were retested in both the ALF and CTF assay to establish reproducibility. Next, to account for error during array generation, plasmids from the 245 locations on the array were sequenced to establish the identity of the over-expressed gene. We observed an error rate of 10% and in these cases these genes were retested for ALF and CTF phenotypes using direct transformations.

**Figure 1.**
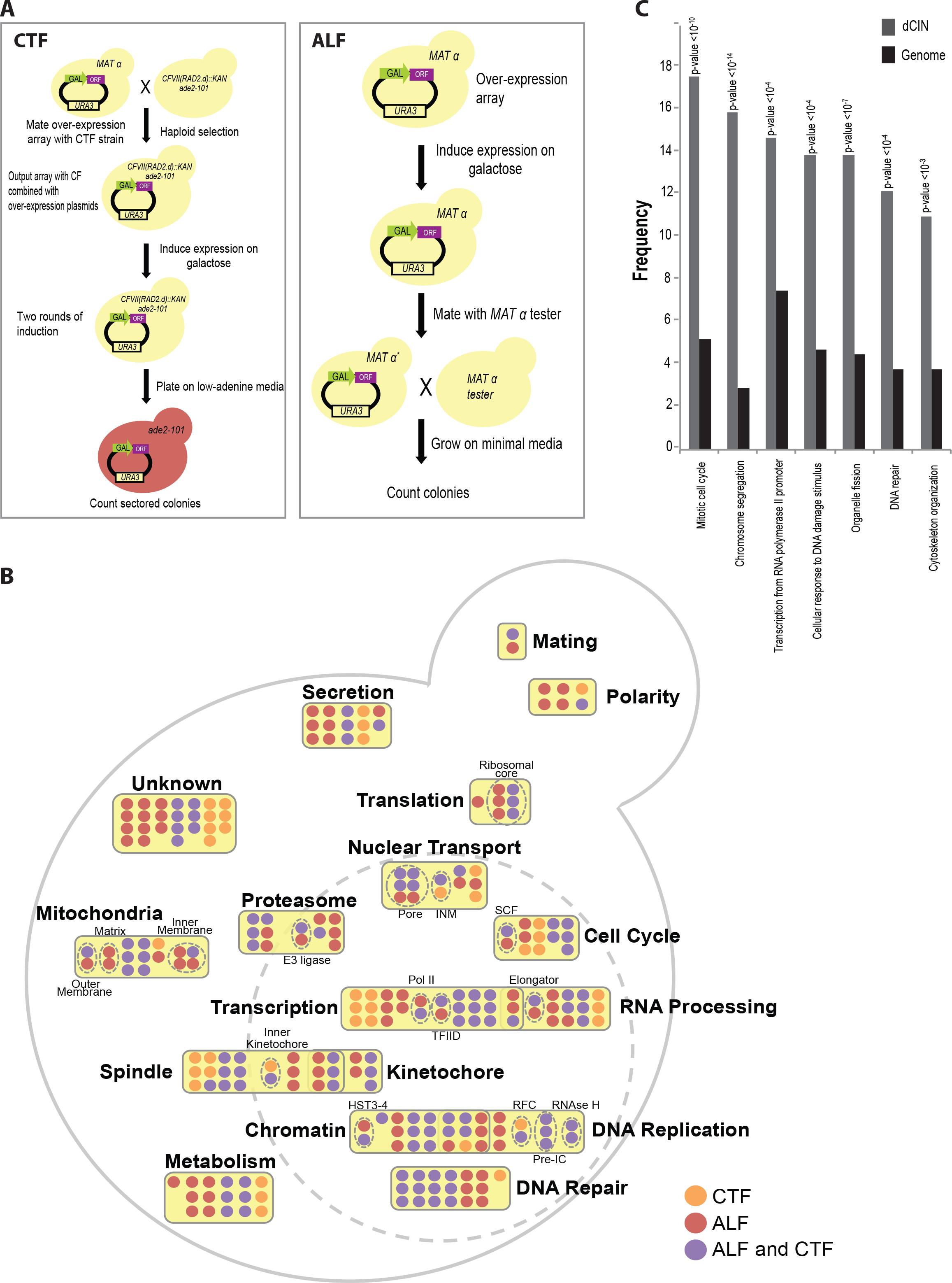
Dosage chromosome instability genes in yeast. (A) Schematics of the genome instability screens. For the CTF screen (left), a query strain containing the *ade2-101* mutation that blocks adenine production thus gives rise to red pigments that is relieved by the presence of *SUP11 (CFVII(RAD2.d)::KAN)* carried on a non-essential fragment of chromosome VII (CF). Upon haploid selection for strains with the CF, *ade2-101* mutation and over-expression plasmids strains were pinned on galactose to induce expression. Strains carrying CF are unpigmented while those that lose the CF were red. The a-like faker (ALF) screen (right) assays the dedifferentiation of the MAT α locus. The over-expression array in a MAT α background was pinned on galactose to induce expression and mated to an α-mating tester strains 2 days later. Growth of diploid mated progeny was assessed on selective media where ability to form diploids reflect loss, deletion or inactivation of the MAT α locus. (B) The 245 dosage CIN genes arranged according to function (Table S1). Colors of the circles indicate the screens genes were identified in: ALF (red), CTF (yellow) and both (purple). The large dotted circle denotes the nucleus. Bold labels and yellow boxes represent groups and the dotted circles indicate protein complexes. Subgroup abbreviations; INM - inner nuclear membrane; SCF - Skp, Cullin, F-box containing complex; Pre-IC - preinitiation complex; RFC - Replication Factor C; Pol II - RNA polymerase II; TFIID - Transcription factor II D. (C) Bar-graph showing enrichment of the dCIN genes for biological process as defined by GO annotations (grey bars) relative to the genome (black bars).

**Figure S1.**
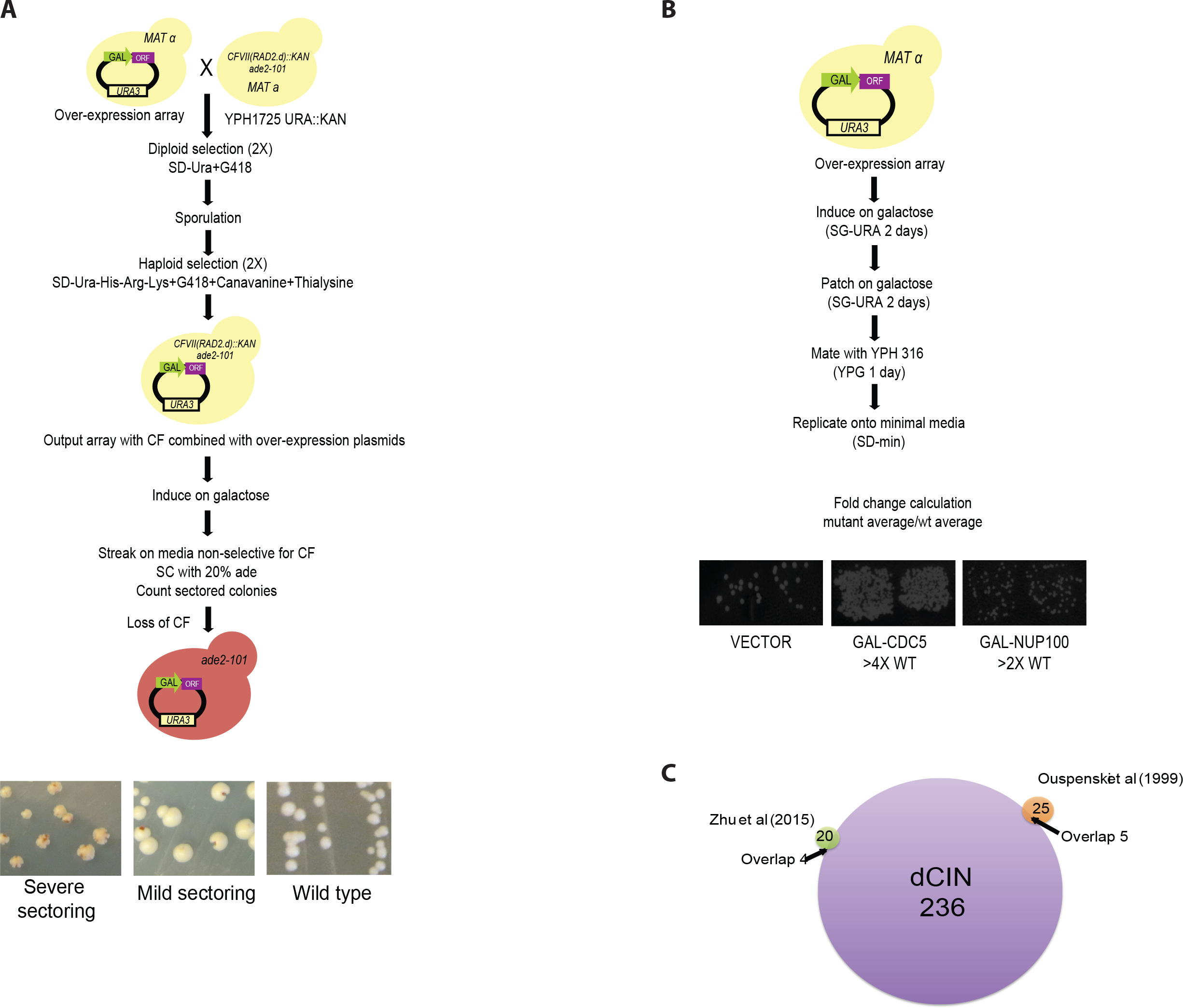
Detailed schematics of the dCIN Screens. (A) Detailed steps involved in the CTF screen. A query strain containing the *ade2-101* mutation that blocks adenine production thus gives rise to red pigments that is relieved by the presence of *SUP11 (CFVII(RAD2.d)::KAN)* carried on a non-essential fragment of chromosome VII (CF). The query was mated to the FLEX array and selected for diploids on selective media. Following sporulation, a haploid output array was generated where each over-expression plasmid was combined with the CTF query. Cells were pinned onto galactose to induce expression and after 48 hours streaked on plates with low adenine media. After growing 7-10 days at room temperature, plates were placed at 4°C for 2-3 days before scoring. Representative images of the qualitative CTF phenotypes are shown below. (B) The a-like faker (ALF) screen assays the dedifferentiation of the MAT α locus. The over-expression array in a MAT α background was pinned on galactose to induce expression and following 48 hours were induced for a second time on galactose plates in 1cmX1cm patches. Cells were mated to an α-mating tester strains 2 days later. Growth of diploid, mated progeny was assessed on minimal media. Representative images from the phenotypes assessed as shown below.

Our screens identified 245 dosage CIN genes (Figure 1B and **Table S1**), enriched for diverse processes including cell division, chromosome segregation, transcription and response to DNA damage (Figure 1C). Over 44% of the genes (108/245) were identified in both ALF and CTF screens, whereas 44 and 93 genes were specific to either the CTF or the ALF screens respectively. These differences reflect the different types of instability detected by the screens. CTF predominantly measures whole chromosome loss where as ALF detects chromosome loss, gene conversion and chromosome rearrangements (Stirling et al., 2011; Yuen et al., 2007). When grouped by biological function based on *Saccharomyces* Genome Database (SGD) and Gene Ontology (GO), a high proportion of dCIN genes associates with nuclear processes and localize to the nucleus. Approximately 30% of the dCIN genes belong to peripheral biological pathways not currently connected to CIN, thus the mechanism of dCIN will require further experiments. Only 24% of the dCIN genes encode for essential proteins showing no bias towards essential genes. The over-expression of 106 dCIN genes are known to decrease cell growth (Douglas et al., 2012), which is not surprising given that high levels of CIN would impact strain fitness.

When combined with previously identified dCIN genes, a total of 290 genes cause chromosome instability when over-expressed (Mishra et al., 2011; Ouspenski et al., 1999; Sarafan-Vasseur et al., 2002; Zhu et al., 2015). Our screens have extended the list of previously known dCIN genes (55) by more than 4-fold, however there is little overlap between the three screens **(Figure 1SC)**.

### Gene deletion and Gene over-expression uncovers distinct chromosome instability genes

Our published CIN dataset includes 692 genes whose reduction-of-function increases genome instability (Stirling et al., 2011). Sixty-seven genes were common to both the CIN and dCIN gene datasets **(Table S2)**. One possibility is that the over-expressed gene acts as an antimorph interfering with the function of the protein or its complex (Prelich, 2012). An explanation for concordance between a gene deletion and the over-expression is disrupting the stoichiometry of a multi-subunit complex, when the removal of a component results in haploinsufficiency (Sopko et al., 2006; Veitia, 2005). Overexpression may titrate shared factors away from a complex or sequester subunits of a complex thereby disrupt multi-protein complexes (Figure 2A). We postulated that if over-expression results in loss-of-function (LOF) we will be able to recapitulated a known negative genetic interaction (GI) between the dCIN gene and an interacting synthetic lethal partner. Negative genetic interactions, such as synthetic sickness (SS) or lethality (SL), take place when the observed fitness of a double mutant is more severe than expected, taking into account the fitness of the two single mutants (Figure 2B; (Mani et al., 2008).

**Figure 2.**
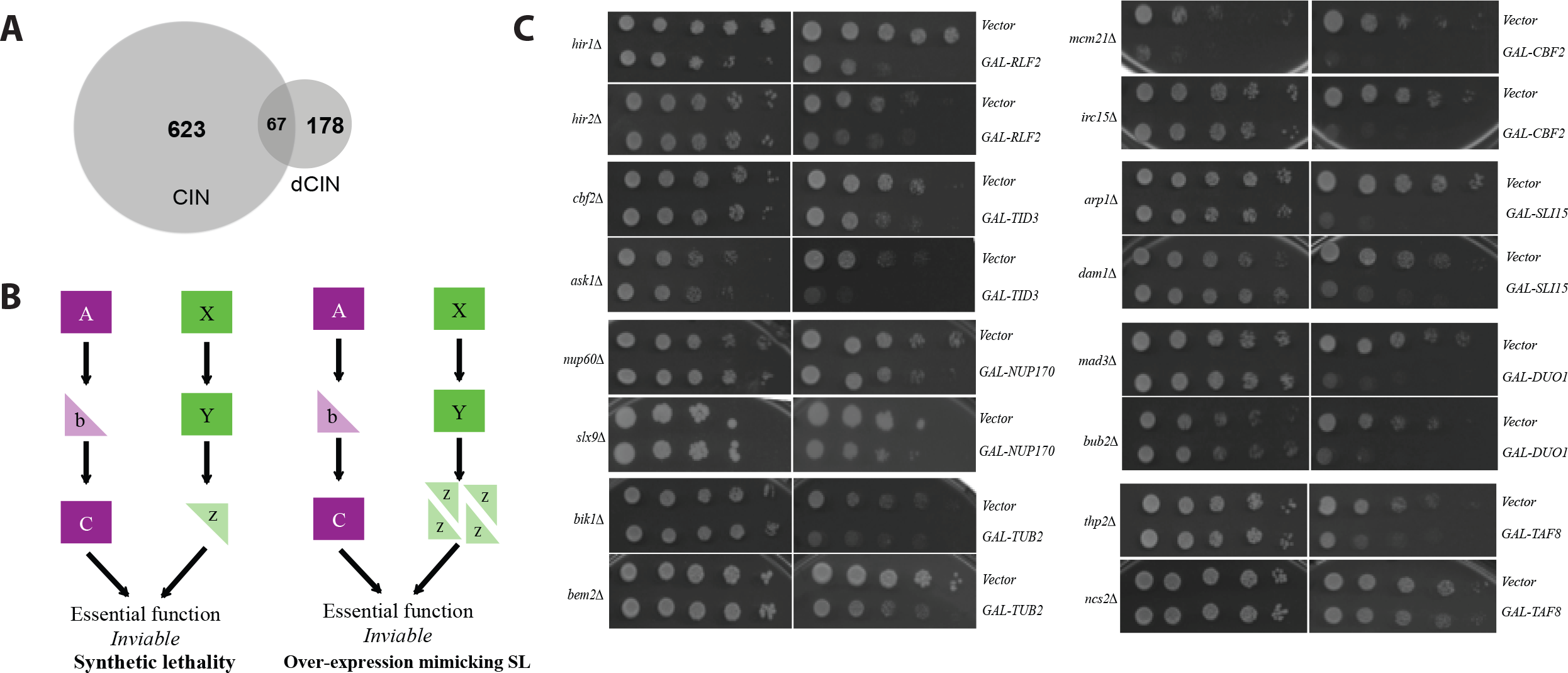
Over-expression recapitulates loss of function. (A) Overlap between the dCIN and the CIN data sets. (B) Negative genetic interactions or synthetic lethality (left) can occur between parallel nonessential pathways when they converge on a common essential biological process. When over-expression results in loss-of-function of a protein (right) disruption of a parallel pathway similarly will lead to synthetic dosage lethality (SDL), recapitulating a negative genetic interaction. (C) Serial spot dilutions of deletion strains transformed with over-expression plasmids and vector control on glucose and galactose containing media.

Thirty-one of the 67 genes encode a component of a protein complex and using the SGD and DRYGIN databases we identified two synthetic lethal partners for each of these genes (Costanzo et al., 2011). Mutant strains were transformed with plasmids to over-express dCIN genes and assayed for a negative GI using serial spot dilutions. Eight genes recapitulated the known GIs suggesting that over-expression may phenocopy LOF (Figure 2C). We also tested the remaining 36 genes, which were not components of protein complexes, using the same assay and 24 recapitulated negative genetic interactions with a known synthetic lethal/sick gene partner **(Table S2)**. A majority of the genes involved in chromosome segregation (10/15) phenocopied LOF. For almost half of these genes (32/67) where both LOF and over-expression results in CIN, therefore the phenotype associated with dCIN is likely due to loss-of-function. *NUP170* is one such example where deletion and over-expression disrupts the nuclear pore (Flemming et al., 2009). In the remaining cases over-expression appears to cause genome instability due to either gain of function or a novel unregulated effect.

### DNA metabolism is affected by gene over-expression

DNA damage and defective DNA repair are well established mechanisms that lead to CIN (Jackson and Bartek, 2009). Therefore we assayed the dCIN genes for the presence of increased DNA damage using Rad52 as a proxy. Rad52 is a protein essential for homologous recombination and form foci in response to double strand breaks (DSBs) and recombination events (Lisby et al., 2001; Symington, 2002). Using SGA we introduced the 245 over-expression plasmids into a reporter strain containing GFP-tagged Rad52 that marked DNA damage sites and mCherry-tagged histone H2A, which marked the nuclei. Spontaneous Rad52-foci were quantified using automated image analysis (Figure 3A and Materials and Methods). Dosage CIN genes that exhibited foci in more than 10% of the cells, a 2-fold elevation above the vector alone controls, from the initial screen were retested. Plasmids were introduced into the reporter strain using direct transformations followed by fluorescence microscopy. After scoring three independent transformants for each dCIN gene, we identified 20 genes whose over-expression increased Rad52-foci (Figure 3B, 3C and **Table S3)**. Majority of the genes identified function in DNA metabolism and chromosome dynamics as expected (Alvaro et al., 2007; Stirling et al., 2012). Four, *CDC6, RRM3, RAD27* and *RAD54*, were previously shown to increase Rad52 foci as LOF mutants (Alvaro et al., 2007; Stirling et al., 2012). We also tested for synthetic dosage lethality interactions (SDL) with *rad52Δ*, since increased Rad52 foci may suggest a requirement for *RAD52* gene function. Dosage CIN genes that induced foci were transformed into a *rad52Δ* mutant strain and were assayed for SDL using serial spot dilutions. Six genes *(AMN1, CDC6, SLK19, PSO2, KAP95, RRM3)* induced SDL, suggesting that strains over-expressing these genes require homologous recombination for viability (Figure 3C).

**Figure 3.**
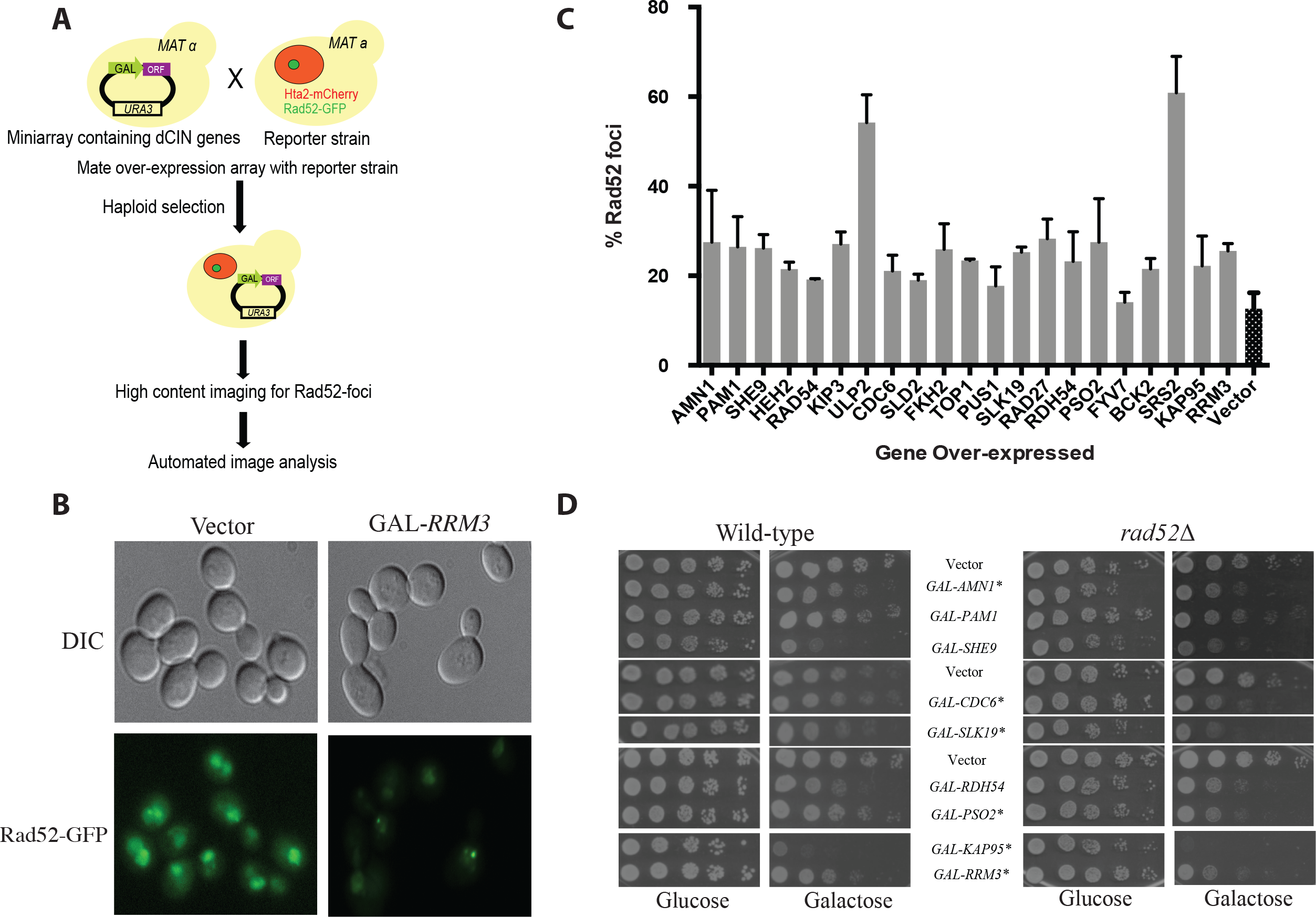
dCIN genes increase DNA damage assessed used Rad52 foci. (A) Schematics of the Rad52-foci screen. A mini array containing the 245 dCIN genes were mated to a reporter strain with Rad52-GFP, to identify DNA damage foci and Hta2-mcherry, which marks the nucleus. Following the generation of a haploid output array, where every over-expression plasmid contained the two reporter genes, gene over-expression was induced by placing the cells on 2% galactose for 16 hours. A high content imaging platform and automated image analysis were used to identify initial hits. (B) Sample images from the Rad52-foci confirmations. The reporter strain transformed with vector alone control show Rad52-GFP localizing to the nucleus. Upon over-expressing *RRM3* in galactose for 16 hours Rad52 forms distinct foci. (C) dCIN genes that show increased Rad52-foci upon over-expression. Each plasmid was directly transformed in to a reporter strains with Rad52-GFP and Hta2-mcherry following image acquisition. Data from three independent experiments, each counting >100 cells are shown. (D) Genetic interactions between the dCIN genes that induce Rad52-foci and cells lacking *RAD52*. Plasmids were directly transformed into either a wild type strain or a rad52Δ and tested using serial spot dilutions. Genes that showed SDL interactions are marked with an (*).

DNA damaging agents are often used to treat tumors that have mutations in genome stability genes (Helleday et al., 2008). Drug sensitivities of dCIN genes may be used for predicting drug sensitivities of tumors with gene amplifications and/or over-expressions. When tested for sensitivity to four genotoxic agents (hydroxyurea (HU), methyl methanesulfonate (MMS), bleomycin and benomyl), 69 dCIN genes were hypersensitive to at least one genotoxic agent **(Table S4)**. One possibility is increased DNA damage that results from gene over-expression may synergize with DNA damaging agents; Pif1 is one such example where over-expression activates the DNA damage response thus inhibiting DNA replication simultaneously with HU leads to cell death (Chang et al., 2009). When over-expression results in LOF the sensitivity profiles will mimic that of the delete. *RTT107* is an example where both the deletion and over-expression results in sensitivity to MMS and bleomycin (Kapitzky et al., 2010). These experiments suggest that these 69 dCIN genes increase chromosome instability by either inducing DNA damage or affecting DNA repair pathways in the cell.

### Over-expressing proteins involved in nuclear-cytoplasmic transport destabilizes the nuclear pore

Genes controlling nuclear-cytoplasmic transport were over represented in our dCIN list (12/47 p-val 2.38e-15) and the nuclear pore complexes play a crucial role in protecting genome integrity (Ibarra and Hetzer, 2015; Rodriguez-Bravo et al., 2014). To deduce the mechanism of dCIN we tested the possibility that over-expression causes defective nuclear pore complexes (NPCs) leading to inadequate transport of proteins involved in essential nuclear processes. We first examined the nuclear localization of green fluorescent protein (GFP) fused to a nuclear localization signal (NLS), to detect defects in general transport. When cells are transformed with a plasmid constitutively expressing NLS-GFP, the fluorescent signal accumulates in the nucleus, whereas defects in transport will change the nuclear accumulation (Shulga, 1996). We detected no observable difference in the distribution of GFP when dCIN genes involved in nuclear-cytoplasmic transport are over-expressed, ruling out general transport defects.

Mutations in Nup170 has a role in chromosome segregation and kinetochore function (Kerscher et al., 2001) and over-expressing the C-terminus of Nup170 causes several nuclear pore proteins to accumulate as cytoplasmic foci (Flemming et al., 2009). Therefore, we next examined the effects on NPC assembly and/or structure by assessing the localization of Pom152, an integral protein of the pore membrane (Wozniak et al., 1994). Over-expressing dCIN genes *HEH2, KAP95, KAP122, NUP53* and *NUP170* caused Pom152-GFP fusion protein to localize into foci (Figure 4). We also tested several additional GFP-tagged NPC components and the distribution of all these components are affected when *HEH2, KAP95, KAP122, NUP53* and *NUP170* are over-expressed (Figure S2). Overproduction of Nup53 generates intranuclear membrane structures with transcisternal pores containing only two integral Nups, Pom152 and Ndc1 (Marelli et al., 2001). From this data we can conclude that a possible mechanism for dCIN induced by genes involved in nuclear-cytoplasmic transport, is through producing defective nuclear pore structures.

**Figure 4.**
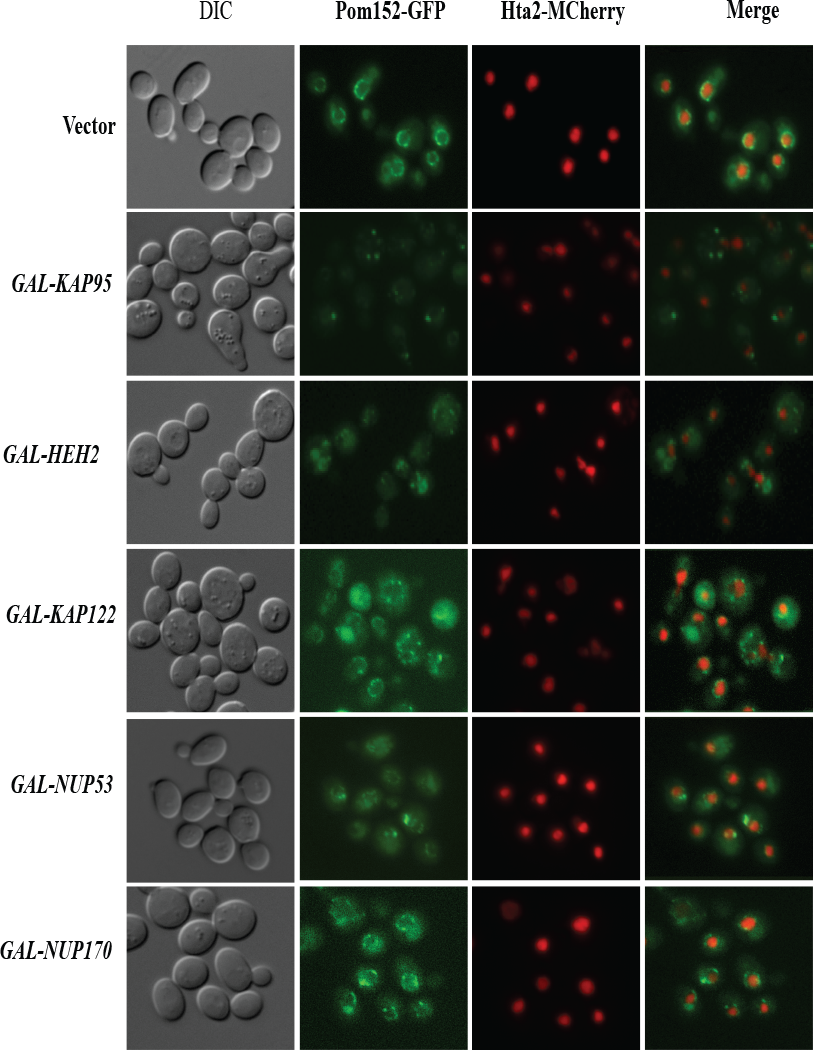
Aberrant localization of nuclear pore proteins upon dCIN gene over-expression. Dosage CIN genes involved in nuclear cytoplasmic transport were directly transformed into a reporter strain with Pom152-GFP, an integral protein of the nuclear pore, and Hta2-mcherry, to mark the nucleus. Over-expressing *HEH2, KAP95, KAP122* and *NUP170* cause Pom152-GFP to accumulate in foci.

**Figure S2.**
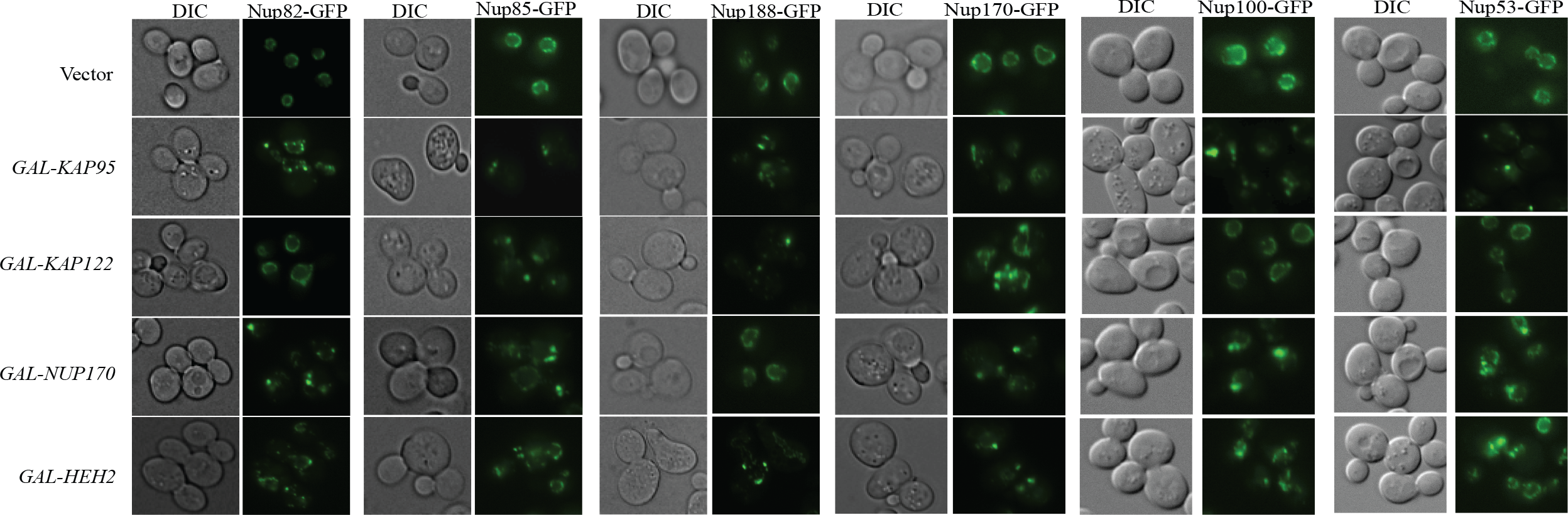
Aberrant localization of nuclear pore proteins upon dCIN gene over-expression. Over-expressing *HEH2, KAP95, KAP122* and *NUP170* cause multiple nuclear pore proteins to accumulate in foci.

### Candidate dCIN genes in humans are amplified and/or over-expressed in cancer

Even with advances made in sequencing technologies defining genetic variants that cause CIN is still rate limiting (Dunham and Fowler, 2013). As majority of the SCNA amplicons contain several candidate genes distinguishing genes that give rise to CIN is equally difficult. We hypothesized similar to previous candidate based studies that identified CIN genes in human cells (Barber et al., 2008; Stirling et al., 2011), the dCIN gene list could be utilized to direct the search for dCIN-associated amplifications in tumors. For this purpose we generated a list of 313 candidate human dCIN genes using sequence homology to yeast dCIN genes (Balakrishnan et al., 2012; O’Brien et al., 2005) **(Table S5)**. There is significant evidence for 8 of these genes where amplification is relevant to the development of cancer (Santarius et al., 2010). Only two genes, *CCNE1* and *CCND1*, have been directly linked to CIN in human cells (Casimiro et al., 2012; Spruck et al., 1999).

Since genome context plays a role on amplification (Gajduskova et al., 2007), we first considered the distribution of human dCIN candidates within frequently amplified or deleted regions in the genome. Thirty-six dCIN genes map to recurrently amplified (19 genes) or deleted (17) regions computed from pan-cancer data **(Table S6)**. *CCNE1* and *CCND1* are two examples of genes within regions frequently amplified in multiple tumor types (Sung et al., 2014). Over-expressing five out of the 19 that map to frequently amplified regions have a role in tumor progression **(see Table S6)**. It is possible that some of the remaining candidate dCIN genes within these frequently amplified regions may be involved in tumor development.

### Over-expressing Tdp1 and Taf12 increase CIN in HT1080 cells

To identify candidate dCIN genes in human cells we chose to use several criteria. These included: (i) increased expression in tumors (FireBrowse Portal), (ii) over-expression causing a biological effect (correlation between expression and clinical outcome) (Davis et al., 2013), (iii) known or predicted roles in tumor progression (oncogene or tumour suppressor) (Davoli et al., 2013; Vogelstein et al., 2013). Our final candidate gene list **(Table S7)** included genes that met at least one of these criteria, to ensure that a broader spectrum of genes will be tested.

We over-expressed candidate dCIN genes in HT1080 cells and examined chromosome instability using a quantitative assay based on a non-essential human artificial chromosome (HAC) (Lee et al., 2013) (Figure 5A). The HAC, which contains *EGFP* (enhanced green fluorescence protein), is maintained as a non-essential chromosome due to the presence of a functional kinetochore. Cells that inherit the HAC fluoresce green while HAC loss will lead to a loss of fluorescence that can be detected using flow cytometry. Gateway-compatible lentiviral expression vectors were used to generate HT1080 HAC-GFP lines over-expressing candidate human genes (Yang et al., 2011). Briefly, full-length genes from the hORFeome collection (8.1) were shuttled into a destination vector that allows high expression levels (CMV promoter) and contained a c-terminal V5 tag. Post-infection expression of the dCIN candidates was confirmed by western blots (Figure S3A). For genes that were not included in the hORFeome collection we choose to test either another component of the complex or a closely related homolog.

**Figure 5.**
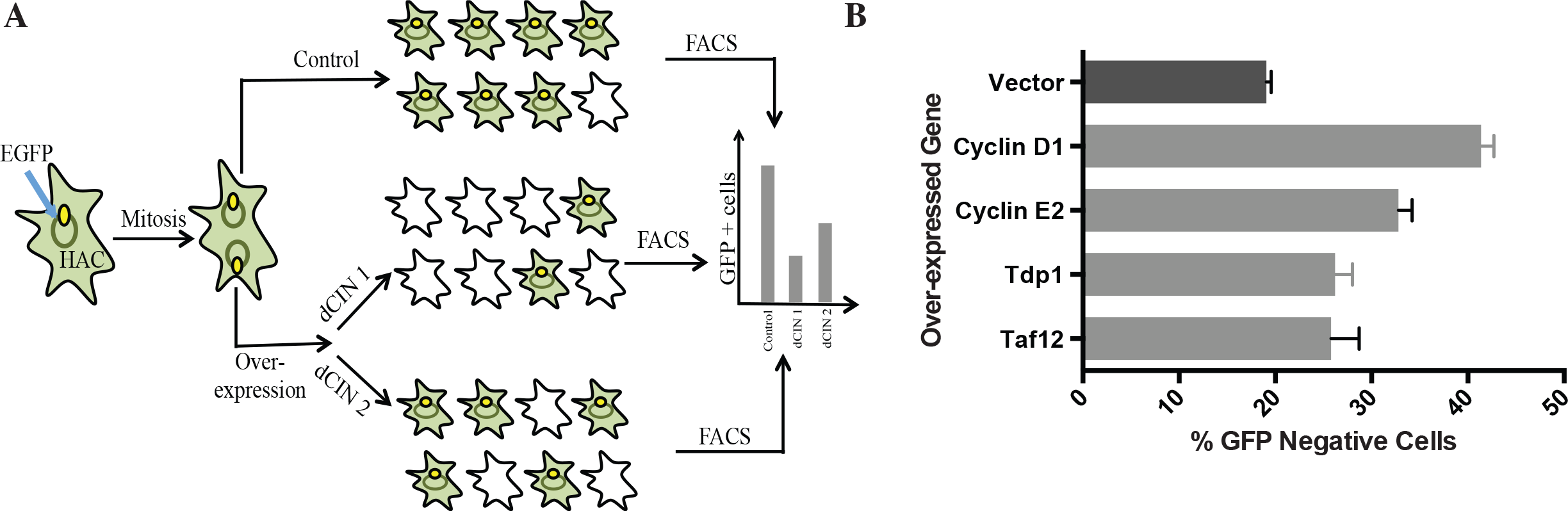
Recapitulating dCIN in human cells. (A) A schematic outlining the human artificial chromosome (HAC) based CIN assay in human cells. Candidate human dCIN genes will be over-expressed in HT1080 containing the HAC-GFP and loss of the HAC, which results in the loss of GFP signal will be quantified using flow cytometry. (B) Over-expressing Cyclin D1, Cyclin E2, Tdp1 and Taf12 induce CIN (light grey bars) compared to HAC-GFP cells over-expressing a vector only (dark grey) control.

**Figure S3.**
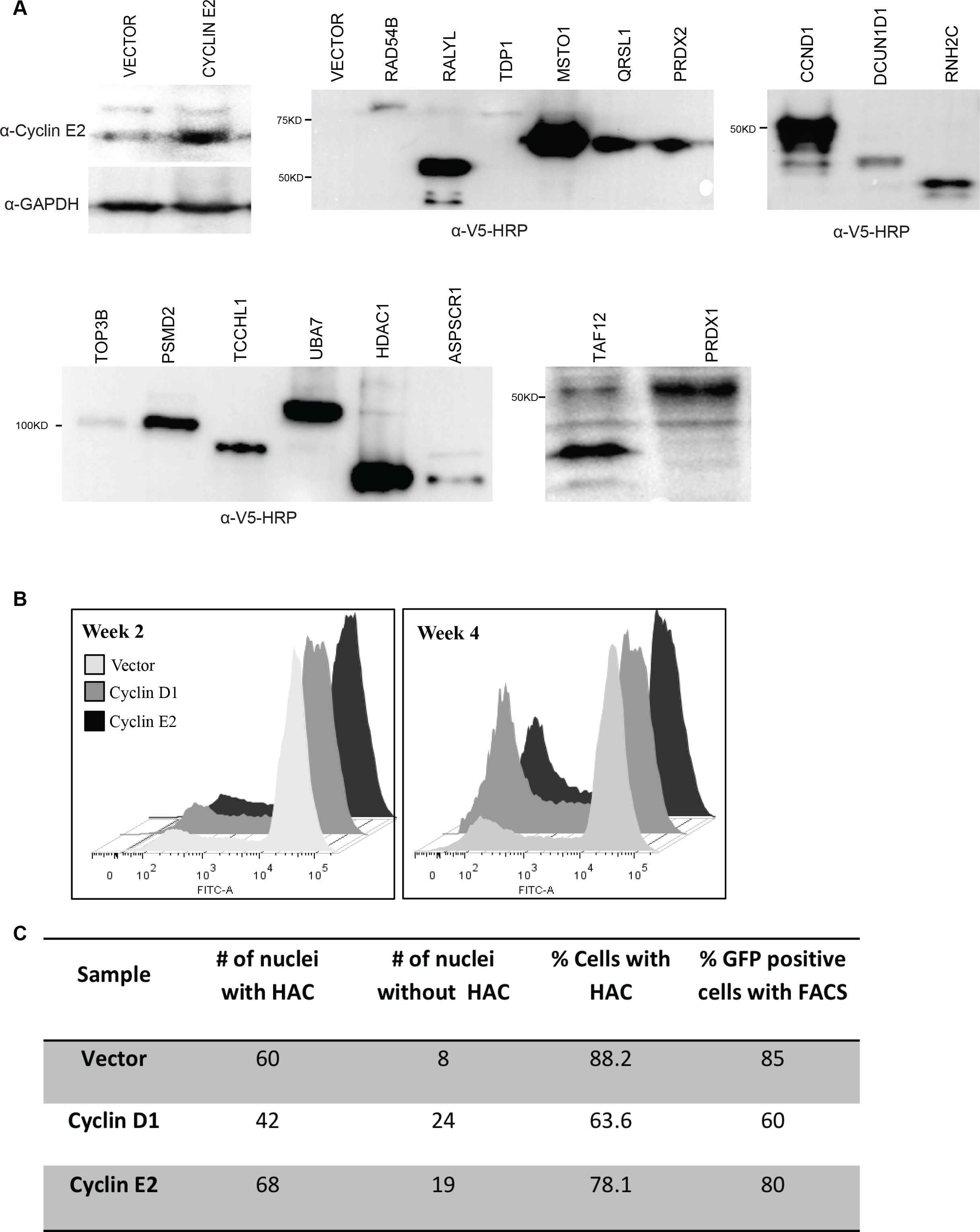
Expression of the candidate dCIN genes tested in human cells. (A) HT1080-HAC lines over-expressing human dCIN candidate genes were assessed for protein expression using the C-terminal V5 tag. (B) Sample plots for HAC-GFP lines over-expressing a vector alone control, Cyclin D1 and Cyclin E2 at 14 days and 28 days following the start of the experiment. Cells with fluorescence values below 10^3^ were considered to be GFP-negative. (C) A table showing FISH data for HAC loss in HAC-GFP lines over-expressing a vector alone control, Cyclin D1 and Cyclin E2 at the end of one experiment.

The HAC-based assay can detect CIN when treated with chemical agents that destabilizes chromosomes (Lee et al., 2013), but had to be tested with genetic perturbations. As a proof of principle experiment, we over-expressed *CCNE2* and *CCND1*, two known human dCIN genes, in HAC-GFP cells. Samples were analyzed using flow cytometry every 7 days for a total of 28 days to determine the proportion of non-fluorescent cells. Over-expression of both Cyclin E2 and Cyclin D1 induced HAC-loss compared to a vector alone control (Figure 6B). As seen by Lee *et al* (2013) there was a delay between HAC loss and the appearance of non-fluorescent cells due to the high half-life of GFP. The difference between the control and cyclin over-expression was clearly distinguishable after 21 days and was more evident after 28 days (Figure S3B). The fraction of cells without the HAC-GFP in the control line coincide with previous studies (Lee et al., 2013). We also confirmed using fluorescent in situ hybridization (FISH) that loss of fluorescence detected by flow cytometry corresponds with HAC loss events (Figure S3C).

**Figure 6.**
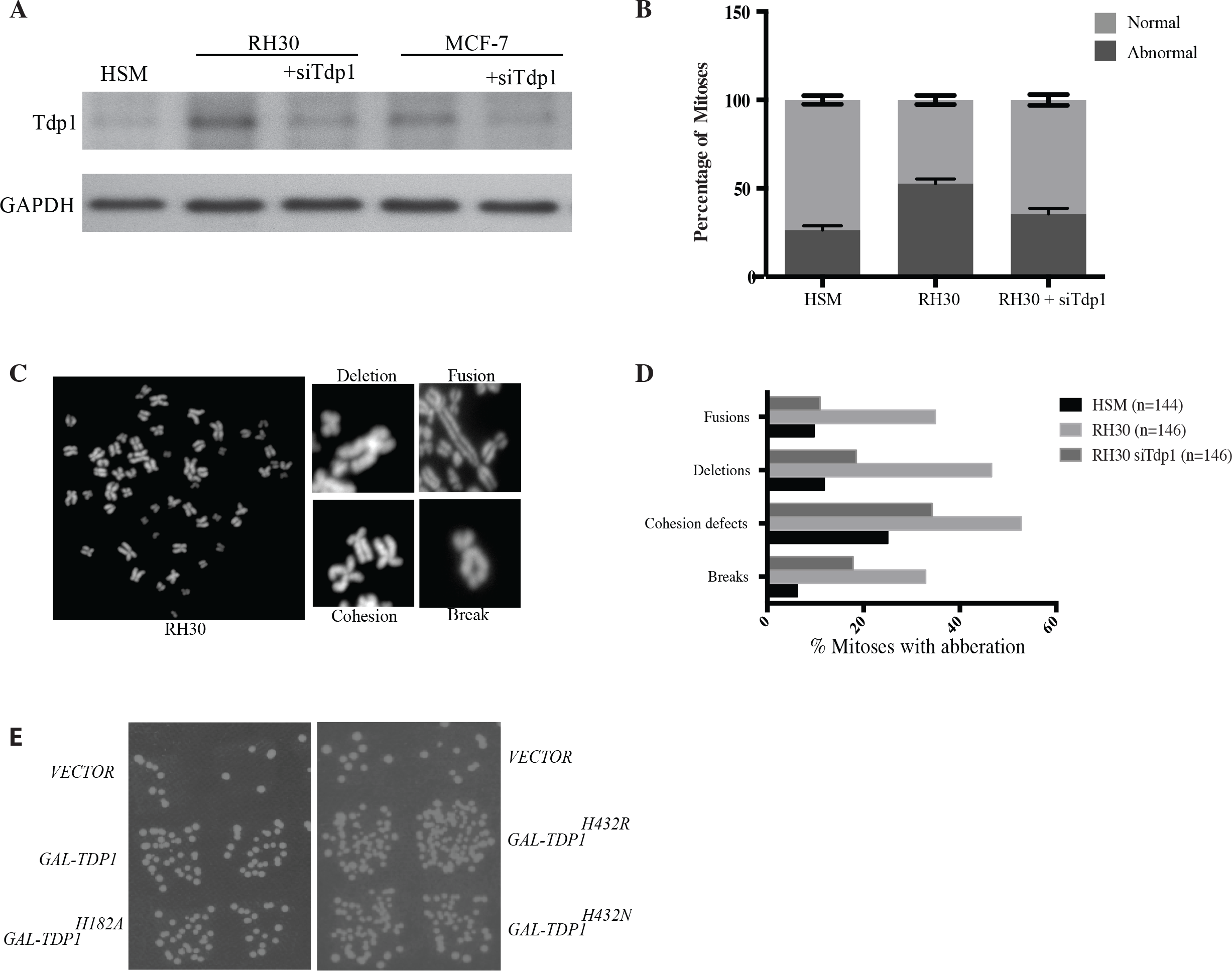
Rhabdomyosarcoma (RMS) cell lines with higher Tdp1 levels show increased CIN. (A) Tdp1 expression levels in RMS cells. Tdp1 knock-down reduces these levels down to wild type levels (HSM). (B) Increased chromosome instability in RH30 cells observed by metaphase chromosome spreads and knock-down of Tdp1 results in the reduction of CIN. Average values from three replicate experiments are shown. (C) Representative chromosome spreads and sample images indicating the types of chromosome abnormalities assessed. (D) Types of chromosome instability detected in RMS cells with high Tdp1 levels. (E) Three catalytic mutants of Tdp1, Tdp1^H182A^, Tdp1^H432R^ and Tdp1^H432N^ were tested in the ALF assay and as expected the phenotype is retained in the Tdp1^H432R^ and Tdp1^H432N^ mutants that are not able to dissociate from the DNA. Catalytic activity of Tdp1, as measured with the Tdp1^H182A^ mutant was not necessary for the ALF phenotype.

Next we tested 18 high priority candidate genes, selected based on the above-mentioned criteria, using the HAC-GFP assay **(Table S7)**. Over-expressing Taf12 and Tdp1 increased the fraction of GFP-negative cells compared to the vector alone control (Figure 6B). Tdp1 is a tyrosyl-DNA-phosphodiesterase with an established role in DNA repair (Yang et al., 1996) and Taf12 is a RNA polymerase II TATA-box binding factor involved in transcription. Taf12 was recently identified as an oncogene in brain tumors (Tong et al., 2015). Increased Tdp1 levels are seen in certain rhabdomyosarcomas (Fam et al., 2013) and non-small cell lung cancer (Liu et al., 2007) and Tdp1 levels also affect the response to chemotherapeutics such as Irinothecan (Barthelmes et al., 2004; Meisenberg et al., 2015). Neither of these genes has been previously linked to CIN in human cells.

**Figure 7.**
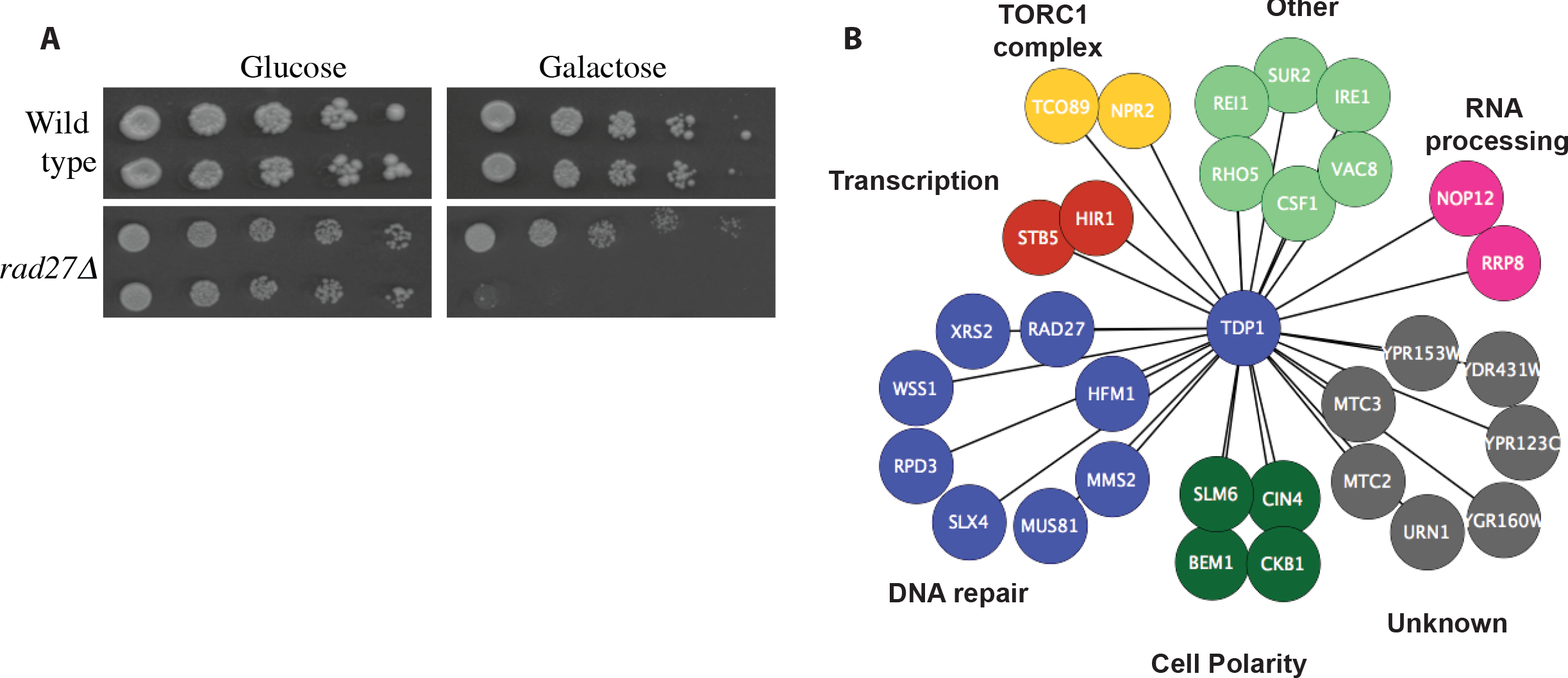
A synthetic dosage lethality screen for Tdp1 identify candidate gene targets for selectively killing Tdp1 over-expressing tumors. (A) A representative image from the TDP1 SDL validations. Over-expressing *TDP1* selectively kill mutants that lack *RAD27*. (B) Synthetic Dosage Lethal (SDL) interactions for yeast Tdp1. Genes are grouped according to biological processes as annotated by Costanzo *et al* (2010a). A complete list of SDL interactions is provided in Table S8.

### Rhabdomyosarcoma lines expressing higher levels of Tdp1 show chromosome instability and catalytic activity of Tdp1 is not necessary for function

Rhabdomyosarcoma (RMS) is one of the most common soft tissue sarcomas in children (Dagher and Helman, 1999). Certain RMS tumors over-express Tdp1 at both the protein and the mRNA level (Fam et al., 2013) and increased levels of Tdp1 have been implicated in resistance of some cancers to Topoisomerase I inhibitor, Irinotecan (Liu et al., 2007; Mascarenhas et al., 2010; Walterhouse et al., 2004). We examined whether increased Tdp1 caused CIN in RMS cells. First, we directly assayed chromosome instability in RH30 RMS cells using metaphase chromosome spreads. RH30 cells express higher than normal levels of Tdp1 (Figure 6A and (Fam et al., 2013)). We see an increase in the number of abnormal mitoses in RH30 cells compared to human skeletal muscle (HSM) cells and this increase in CIN can be abrogated by the knock-down of Tdp1 (Figure 6B). Chromosome defects observed include chromosome fusions, breaks, deletions and cohesion defects all of which are increased in RH30 cells (Figure 6C and 6D).

Next, to determine whether catalytic activity of Tdp1 was necessary for the dCIN phenotype, we used site directed mutagenesis and generated three mutants, Tdp1^H182A^, Tdp1^H432R^ and Tdp1^H432N^. 182 432 182 Both H^182^ and H^432^ are located in the active site of Tdp1 and mutating H^182^ to an alanine, greatly reduces the activity of Tdp1 (Interthal et al., 2001; Liu et al., 2007). H^432^ is necessary to dissociate Tdp1 from the DNA and this residue (H^493^ in humans and H^432^ in yeast) is mutated to an arginine in patients with a rare recessive neurodegenerative disease spinocerebellar ataxia with axonal neuropathy (SCAN1) (Takashima et al., 2002). A H432N mutation causes severe growth defects in yeast (He et al., 2007). Over-expressing both Tdp1^H432R^ and Tdp1^H432N^ mutants further increased CIN using the ALF assay, which was expected since these mutants are unable to hydrolyze the Tdp1-DNA links (Figure 6E). However, there was no reduction in CIN when the catalytically inactive Tdp1^H182A^ mutant was over-expressed as we expected. This observation is supported by recent findings where Tdp1^H182A^ retains catalytic activity *in vivo* (Comeaux et al., 2015). This experiment suggests that it is not the increased enzymatic activity of Tdp1 that is responsible for the dCIN phenotypes. One possibility is that increase amounts of Tdp1 will promote binding to the DNA non-specifically, thereby introducing DNA adducts to the cells. In a manner similar to Top1-DNA adducts these may lead to increased DNA damage and CIN (Liu et al., 2002).

### A synthetic dosage lethality screen for Tdp1 in yeast to identify candidate targets for specifically targeting tumors with increased Tdp1 levels

To identify candidate drug targets for tumors with high Tdp1 levels, such as Rhabdomyosarcoma, we performed a genome-wide synthetic dosage lethality screen in yeast. Briefly, we introduced a query strain over-expressing Tdp1 into the deletion mutant array using SGA (Winzeler et al., 1999). The haploid output array where the deletions were combined with the over-expression were placed on media containing galactose to induce over-expression, followed by assessing colony size as a proxy for fitness (Douglas et al., 2012; Sopko et al., 2006). After validating our initial SDL interactors with direct transformations, we identified 31 genes that caused SDL upon Tdp1 over-expression (Figure 7A, 7B and **Table S8)**. Our screen was enriched for genes involved in response to DNA damage (p-value = 2.15x10^-05^), which is expected given the role of Tdp1 in DNA repair. Twenty three of these genes have known human homologues (Balakrishnan et al., 2012).

Several of these candidate genes such as *RAD27, WSS1, SLX4* and *MUS81* have physical, genetic or functional interactions with Tdp1. Rad27 and Tdp1 are both involved in the repair of covalent Top1 complexes and removal of Rad27 renders cells slightly sensitive to camptothecin, a topoisomerase inhibitor (Deng, 2005; Vance and Wilson, 2002). Wss1, a metalloprotease that functions in DNA repair (Biggins et al., 2001), has been shown to be essential for viability following topoisomerase I-mediated DNA damage (Stingele et al., 2014). Both *SLX4* and *MUS81*, endonucleases are involved in replication associated DNA repair (Fricke and Brill, 2003; Interthal et al., 2001) and protect cells against camptothecin toxicity (Deng, 2005). Therefore it is possible that one common mechanism for SDL between these genes and Tdp1 may relate to high levels of Tdp1 interacting non-specifically with the DNA, a condition that makes all these genes that protects the genome against DNA-protein adducts essential to these cells.

## Discussion

Previous studies in budding yeast uncovering loss-of-function (Yuen et al., 2007) and reduction-of-function (Stirling et al., 2011) alleles of CIN genes have been instrumental in identifying and characterizing chromosome instability genes in humans (Barber et al., 2008; McManus et al., 2009). We chose to complement this list with gene over-expression to assay gain-of-function phenotypes. Here, we report two genome-wide screens for dosage CIN (dCIN) genes in yeast, genes whose over-expression increases chromosome instability. The overlap between our screens and the previous dCIN screens is low, where we only identified ~15% of the genes identified in either screen, however it is better than the lack of any overlap between the two previous screens (Ouspenski et al., 1999; Zhu et al., 2015). These differences could be due to the differences in the chosen method for gene over-expression, galactose induction in Ouspenski *et al* and this study versus the MoBY-ORF collection (Ho et al., 2009) where genes are expressed from the endogenous promoter in Zhu *et al*. Also, the use of multiple CIN assays rather than just the CFT assay we uncovered 93 additional genes that were only identified in the a-like faker screen.

With these screens we have now a comprehensive collection of CIN genes in yeast that includes loss-of-function, reduction-of-function and over-expression. The complete catalogue amounts to 900 genes, 692 CIN genes and 292 dCIN genes, which are necessary for maintaining genome stability. While our dataset is enriched for processes known to be involved in genome instability, DNA damage repair, chromosome segregation, cell cycle phase and transcription (Skoneczna et al., 2015), we also identified genes in pathways that have only recently been linked to genome instability such as metabolism (Shaukat et al., 2015) as well as processes not directly involved in genome instability such as secretion, suggesting that we have yet to fully understand the role chromosome instability plays in human disease. The breadth of this collection also highlights the significance of chromosome instability particularly in cancer progression, thus has important practical implications.

A third of the genes that caused CIN when over-expressed, also caused CIN when their function was perturbed, and 46% of these over-expression constructs also recapitulated known synthetic lethal phenotypes for the reduction of function alleles suggesting that over-expression in these cases might phenocopy a reduction of function possibly due to perturbation of protein complex stoichiometry (Veitia, 2005)., However, we did not find an association between genes that phenocopied LOF and proteins that are annotated to be components of complexes, in agreement with previous studies (Sopko et al., 2006). Genes involved in regulating the cell cycle and chromosome segregation were overrepresented in the subset of genes in which over-expression phenocopied deletion. These essential processes are more sensitive to changes in protein levels, since perturbing the dosage of genes involved in the mitotic cell cycle and DNA processing, through over-expression or deletion, affects cell growth negatively (Yoshikawa et al., 2011). Therefore, in these cases disruption of an essential process by either over-expression or LOF may explain the concurrence in the CIN phenotype. Our work with genes involved in nuclear-cytoplasmic transport also lends support to this conclusion. As expected, perturbing protein levels of nuclear pore components via deletion or over-expression resulted in structural alterations to the nuclear pore (Flemming et al., 2009). For the remaining gene, where there was no phenotypic concurrence, it is possible that over-expression results in a gain-of-function phenotype (Sopko et al., 2006), however direct test will be necessary to determine this possibility.

Pathways involved in DNA metabolism are integral to genome maintenance and the over-expression of more than a third of dCIN genes either increased DNA damage (measured by increased Rad52 foci) or were sensitive to DNA damaging agents. Pif1 is the only previously identified gene whose over-expression increases sensitivity to DDAs (Chang et al., 2009). Earlier cell biological screens have looked at the effects of LOF (Alvaro et al., 2007) and reduction-of-function alleles (Stirling et al., 2012) involved in alteration to HR pathways, however, our study is the first to look at effects of over expression on Rad52-foci formation. While a majority of these genes belong to biological pathways previously linked to DNA metabolism (Alvaro et al., 2007), we identified several genes with no previous links to DNA damage or sensitivity to DDAs, such as *PAM1, SHE9, PUS1* and *KIP3*. Both *KIP3* and *SHE9* show genetic interactions with genes involved in DNA metabolism (Costanzo et al., 2011; Dixon et al., 2008) and Pam1 physically interacts with another dCIN gene Cmr1, a DNA binding protein involved in repair (Gilmore et al., 2012) offering possible links to DNA metabolism. The role of *PUS1*, a pseudouridine synthase that is conserved in humans (Carlile et al., 2014), in DNA damage remains to be seen, suggesting that a genome-wide over-expression screen for Rad52-foci may identify additional genes involved in DNA metabolism. Approximately 65% of the dCIN genes that showed sensitivity to DNA damaging agent(s) have human homologs, thus it may be possible to leverage these sensitivity profiles to predict drug sensitivities of tumors that are specifically over-expressing these homologs. This dataset thus provides a rich source of information to extend the existing practice of using DNA damaging agents to treat tumors with defects in genome stability genes (Helleday et al., 2008).

To determine whether dCIN genes can be used to identify dCIN genes in humans and to uncover amplified and/or over-expressed genes relevant to tumor progression, we generated a list of human homologues of yeast dCIN genes. In addition to two previously characterized dCIN genes in humans (Caldon et al., 2013; Casimiro et al., 2012; Spruck et al., 1999), our dCIN list captured 8 genes whose over-expression had a clear role in cancer progression (Santarius et al., 2010), as well as genes that mapped to frequently amplified and/or deleted regions in cancer cells. However, when directly tested 20 candidate genes for dCIN in human cells using our HAC-based assay we only identified 2 new dCIN genes in human cells. There are several possible reasons for this observation. It is possible that the remaining 16 candidate genes do not contribute to chromosome instability in human cells. Similar to the CTF assay in yeast the HAC-based CIN assay requires the complete loss of the human artificial chromosome for detecting CIN (Lee et al., 2013). For this reason multiple CIN assays are utilized in yeast to uncover CIN genes and many of these genes that are specific to a single assay (ALF or CTF for example) signify the differences in mechanisms used to generate instability. Therefore, it is possible that the sensitivity of our assay is too low to distinguish genes that cause subtle CIN phenotypes. The stability of GFP and the length of time an experiment runs (4 weeks) are two other caveats of this CIN assay, however, when compared to other CIN assays such as metaphase chromosome spreads and micronuclei assay, it is less labor-intensive and more reproducible even between laboratories (Crasta et al., 2012; Lee et al., 2013). Finally, because we used the criteria of either amplification or over-expression in order to capture a broad spectrum of candidates to be tested, our candidate gene list included genes that were amplified but not over-expressed in tumors, which may have also contributed to the lower numbers of dCIN genes confirmed in human cells.

*TDP1* and *TAF12* the two novel dosage CIN genes in humans identified in this study, to our knowledge have not previously been linked to chromosome instability. *TAF12*, transcription initiation factor IID subunit 12, is a component of the TFIID general transcription factor that is involved in RNA polymerase II-dependent transcription (Mengus et al., 1995). Recent work identified *TAF12* as an oncogene involved in the formation of chroid plexus carcinomas, a frequently lethal brain tumors (Tong et al., 2015). Transcription is a process that has many links to genome instability and aneuploidy-induced transcriptional changes is just one possible method employed by cancer cells to escape cellular homeostasis mechanisms (Giam and Rancati, 2015). Tdp1 expression is increased in several specific types of cancers including non-small cell lung cancers (Liu et al., 2007), Dukes ‘C colorectal cancers (Yu, 2005) and some Rhabdomyosarcomas (RMS) (Fam et al., 2013). We specifically selected Tdp1 to be tested as a dCIN candidate in human cells because it was over-expressed in RH30 Rhabdomyosarcoma cells (Fam et al., 2013). We show that high levels of CIN in RH30 cells can partially rescued by the knockdown of Tdp1, suggesting that Tdp1 is indeed responsible for the CIN phenotype. This also implies that the high chromosome instability seen in certain subtypes of Rhabdomyosarcomas (Goldstein et al., 2006) may be due to Tdp1. The failure of clinical trials involving Irinotecan has been attributed to increased levels of Tdp1(Hosoi, 2015; Mascarenhas et al., 2010; Pappo et al., 2007) it is possible that in addition to Tdp1 levels, the chromosome instability induced by Tdp1 may be an additional mechanism employed by cancer cells to develop resistance to chemotherapeutic agents. Alternatively, Irinotecan may not be effective since increased Tdp1 levels are already forming non-specific adducts on the DNA making Irinotecan ineffective, because Tdp1 blocks Top1 from accessing the DNA. This model is supported by the yeast data where catalytic activity of Tdp1 is not necessary for the CIN phenotype. The increase in CIN observed when Tdp1^H432R^ and Tdp1^H432N^ mutants are over-expressed further supports this idea, since these mutants are unable to dissociate from the DNA (Takashima et al., 2002).

Our *TDP1* SDL screen aimed to identify targets, where higher levels of Tdp1 will render the cells more susceptible to the knockdown of a second gene and our screen was enriched for genes involved in DNA metabolism, as expected. Given that Tdp1 catalytic activity is not necessary for its CIN phenotype, it is possible that when Tdp1 is over-expressed it bind to DNA non-specifically blocking other repair proteins from accessing the DNA. In this model, as with trapped Top1-adducts on DNA upon treatment with Top-1 poisons, enzymes involved in removing protein-DNA adducts become necessary to remove excess Tdp1-DNA adducts and to maintain cell viability. This explains, why the removal of several proteins that function in parallel or together with Tdp1 such as *WSS1, RAD27, SLX4* and *MUS81* is synthetic dosage lethal with *TDP1*.

We present here a platform that can be leveraged to identify and characterize dosage chromosome instability genes in cancer. Dosage chromosome instability genes identified in the budding yeast can provide valuable insight into identifying and characterizing chromosome instability genes in humans. This dataset may also provide insight into over-expressed genes within tumors that contribute to tumor progression, partly addressing the challenge of uncovering driver genes within SCNAs (Davoli et al., 2013; Zack et al., 2013). Genetic context of cancer cells often will define their sensitivity profiles to chemotherapeutics (Hart et al., 2015) and screens in well characterized cancer cells will tell us which of these genetic vulnerabilities can be exploited in a given tumor type (Boehm and Golub, 2015). We propose synthetic dosage lethality screens as a powerful approach for fast and easy identification of chemotherapeutic drug combinations within the context of over-expression that will enable us to specifically target these cells. Combining dosage CIN with synthetic dosage lethality thus offers the potential to expand the chemotherapeutic space that could be utilized to target cancer cell vulnerabilities.

## Materials and Methods

### Yeast Strains, Growth Conditions and Plasmids

The *S. cerevisiae* strains and plasmids used are listed in **Table S9**. Standard methods and media were used for yeast growth and transformations. The expression of genes under the *GAL1* promoter in the liquid growth assays were induced for 16 hours by adding 2% galactose to synthetic media lacking uracil. Synthetic minimal medium supplemented with appropriate amino acids was used for strains containing plasmids. Site-directed mutagenesis of *MPH1* and *UBP12* in pDONR221 was performed using a QuickChange^TM^ kit (Stratagene) following the manufacturer’s protocols. All clones were confirmed by sequencing.

### Dosage CIN Screens and confirmations

**CTF Screen:** SGA screens were performed to introduce the CTF markers *(ade2-101*::NatMX and CFVII *{KANMX, SUP11})* to the FLEX over-expression array (Douglas et al., 2012), using a Singer RoToR as described in (Tong et al., 2001). After diploid selection and sporulation, haploids that are Ura+, Kan^R^ and Nat^R^ were selected. Gene expression was induced on galactose for 2 days and tested for stability of the CF as in (Spencer et al., 1990). Candidate dosage CIN genes identified in the genome-wide screen were confirmed using direct transformations. Three independent transformants were tested for CTF and were assessed qualitatively by eye.

**a-like Faker Screen:** Screen was performed as highlighted in Warren *et al* (2003). Over-expression was induced for 2 days, each strain was patched out on galactose in 1-cm2 patches and mated to a *Mat a his1* tester lawn by replica plating on galactose containing media. His+ prototrophs were selected on synthetic complete medium lacking histidine, uracil, lysine, adenine, tryptophan, and leucine. Candidate dosage CIN genes identified in the genome-wide screen were confirmed using direct transformations with three independent transformants. Colonies on final plates were counted and ratios of vector to over-expressed gene were calculated. Genes whose over-expression that cause >2-fold increase in chromosome instability is reported in Table S1.

### Sensitivity to Genotoxic agents

Each dCIN candidate was grown in liquid media containing glucose followed by serial spot dilutions. Spot assays were quantified by eye to detect a difference in colony size observed on genotoxic agents.

Final concentrations of genotoxic agents on plates: hydroxyurea (Sigma; 100mM), MMS (Sigma; 0.001%), bleomycin (Enzo life Sciences; 10ug/ml) and benomyl (Sigma; 10ug/ml).

### SDL Screens and Confirmations

A query strain with a plasmid containing GAL*-TDP1* was crossed to the yeast deletion array using SGA (Tong et al., 2001). Using a series of replica pinning steps an output array was generated where each deletion mutation on the array was combined with the plasmid for over-expressing *TDP1*. Over-expression of Tdp1 was induced by pinning on to media containing galactose, followed by data analysis as previously described (McLellan et al., 2012). All interactions that met a cut-off of >25% changes in growth when compared to a vector alone control were chosen for validation. For confirmations each deletion strain was transformed with plasmids containing a vector alone control or *GAL-TDP1*. Transformants were tested using serial spot dilutions and only the confirmed interactions are reported in Figure 7.

### Cell Biology

#### Rad52-foci Screen and confirmations

A query strain containing endogenously expressing Rad52-GFP and Hta2-mCherry (genotype: MATa Rad52-GFP::KanMX, Hta2-mCherry::NatMX, his3Δ1, leu2Δ0, ura3Δ0) was introduced to a mini array consisting of 245 FLEX array plasmids (Hu et al., 2007) using SGA (Tong et al., 2001). Haploid strains derived from SGA were inoculated into liquid medium (S_Raffinose_ - uracil - histidine + NAT + G418) and grown to saturation overnight. Saturated cultures were then diluted and grown to mid-log phase in low fluorescent medium (LFRaffinose - uracil - histidine + NAT + G418). Cell cultures were induced with 2% final concentration galactose for six hours at 30°C. A measurement of OD_600_ was used to identify an ideal concentration of cells to image, and this volume of cell culture was transferred to PerkinElmer CellCarrier plates (PerkinElmer 6007558), and subsequently imaged using an Evotec Opera high-throughput confocal microscope with a 60X objective (PerkinElmer). Single plane confocal images of live yeast cells were recorded from four positions per well. Images from the GFP and RFP were acquired simultaneously to avoid cell movement during imaging. CellProfiler^TM^ version 1.0.5811 (Carpenter et al., 2006) was used to detect cells and nuclei in yeast confocal images using segmentation-based approaches. Intensity, texture and morphological measurements were extracted from each identified object. CellProfiler^TM^ can be downloaded from http://www.cellprofiler.org/. Classification was used to detect DNA damage foci in cellular objects identified and measured using CellProfiler^TM^ image analysis software (Styles and Andrews, unpublished data).

Genes that resulted in >10% spontaneous foci in the array-based screen were confirmed using retransformation. Cells were grown to saturation in glucose media and shifted to galactose for 16 hours before imaging. Cells were imaged using Zeiss Zxioscope and analyzed using Metamorph (Molecular Devices). Over 300 cells were counted for each over-expressor.

#### Nuclear pore imaging

A wild type strain was co-transformed with a plasmid constitutively expressing NLS-GFP which accumulates in the nucleus of growing cells (Shulga, 1996) and plasmids to over-express NPC genes and imaged as above. Localization of NLS-GFP were quantified by defining pixel area within the nucleus and another, equally sized area immediately adjacent in the cytosol as reported previously (Minaker et al., 2013). GFP-tagged c-terminal proteins from the yeast GFP collection (Huh et al., 2003) were introduced to a strains with Hta2-mcherry to mark the nucleus. These strains were transformed with plasmids over-expressing NPC genes and imaged as above.

### Cell Culture

Human fibrosarcoma (HT1080) cells with alphoid^tetO^-HAC-GFP were cultured in Dulbecco’s modified Eagle’s medium (DMEM) (Life Technologies) with 10% FBS. To maintain the HAC, HT1080 cells were grown in 5 μg/ml of blasticidin (Sigma) as previously described (Lee et al., 2013). RH30 (alveolar rhabdomyosarcoma, PAX3-FOXO1) cells were cultured in Dulbecco’s modified Eagle medium (DMEM; Gibco BRL Life Technologies) supplemented with 10% heat-inactivated FBS (Hyclone) and 1% antibiotic-antimycotic (Gibco BRL Life Technologies). Human skeletal myoblast cells were cultured using skBM-2 (Lonza) supplemented with 15% FBS, 1% antibiotic-antimycotic and 9 g/L D-glucose (final concentration 10 g/L glucose). Rhabdomyosarcoma cell lines were procured from the Leibniz Institute DSMZ-German Collection of Microorganisms and Cell Cultures or the American Type Culture Collection. The human skeletal muscle cell-line was obtained from Lonza (XM13A1). All cells were grown at 37°C with 5% CO2 in a humidified environment.

### Gene over-expression using lenti-viral vectors

Clones for candidate human genes were taken from the hORFeome 8.1 collection, sequenced and shuttled into the destination vector pLX302 PURO DEST (Addgene plasmid no. 25896). Lentivirus was made in HEK293T cells using MISSION lentiviral packaging mix (Sigma) and Fugene 6 (Promega) optimized according to manufacturer’s instructions.

### HAC-GFP assay

HAC-GFP cells over-expressing the dCIN candidates were grown without selecting for the HAC (blasticidin). Cells were passaged as required (2-3 days) and samples were examined every 7 days for loss of GFP signal using flow cytometry.

### Flow cytometry

Cells were harvested using trypsin treatment, resuspended in PBS (phosphate buffered saline) and fluorescence was determined using flow cytometry. EGFP expression was detected using a FACS Calibur (BD Bioscienes) using FACS DIVA software (BD Bioscienes) and were analyzed using FlowJo (FlowJo, LLC). A minimum of 4×10^4^ events was acquired for each sample.

### FISH analysis with the PNA probe

The presence of HAC in an autonomous form was confirmed by FISH analysis as previously described (Lee et al., 2013).

### Immunobloting

Cells were collected after trypsinization and centrifugation. Pellets were resuspended in 50mM Tris-HCL (pH 7.5), 150mM NaCl, 10% glycerol, 1% Triton X-100, and protease inhibitors (Roche), followed by lysis by sonication and centrifugation to remove debris. Lysates separated by SDS-PAGE, transferred to PVDF were blotted with either a monoclonal V5 antibody (Invitrogen; R96025) or a HRP-conjugated V5 antibody (Invitrogen; R96125).

### Tdp1 siRNA knock-down experiments

*TDP1* was knocked down transiently by transfecting 8 x 10^4^ cells with 100 nmol/L of pooled siRNAs (Dharmacon) in a 24-well plate using Lipofectamine 2000 (Invitrogen). Knockdown was confirmed by immunoblotting and qRT-PCR. The sequences of the siRNAs are listed in **Table S9**.

### Chromosome spreads

Mitotic chromosome spreads were performed as described in (Barber et al., 2008). Actively growing cells were treated with 0.1μg/ml colcemid for 2 hours before washing, trypsinization and adding hypotonic (0.075M KCl) for 5 minutes at room temperature. Cells were fixed three times with 3:1 methanol:glacial acetic acid, centrifuged (800rpm, 5 miutes) before mounting on clean slides. Slides were stained with DAPI before acquiring images with a Zeiss Axiovert 200 microscope, a Zeiss AxiocamHR camera, and the Zeiss Axiovision imaging system. Two independent observers assessed chromosome abnormalities in each of the images.

## Acknowledgements

We thank you Dr. Nigel O’Neil and Dr. Peter C. Stirling for their helpful discussion and comments. This work was supported by grants from the Canadian Institute of Health Research (MOP 38096) and the National Institute of Health (RO1CA158162) to P.H. S.D. was supported by a Canadian Institute for Advance Research Global Fellowship and a Canadian Institute for Health Research Banting Postdoctoral Fellowship. P.H. and B.J.A are Senior Fellows in the Genetics Networks program at the Canadian Institute for Advanced Research. The authors declare no conflict of interest.

